# Translocation of flexible and tensioned ssDNA through *in silico* designed hydrophobic nanopores with two constrictions

**DOI:** 10.1101/2020.05.15.097485

**Authors:** Punam Rattu, Taylor Haynes, E. Jayne Wallace, Syma Khalid

**Affiliations:** School of Chemistry, University of Southampton, Highfield Campus, Southampton, SO17 1BJ; Oxford Nanopore Technologies Ltd, Oxford

## Abstract

Protein-inspired nanopores with hydrophobic constriction regions have previously been shown to offer some promise for DNA sequencing. Here we explore a series of pores with two hydrophobic constrictions. The impact of nanopore radius, the nature of residues that define the constriction region and the flexibility of the ssDNA is explored. Our results show that aromatic residues slow down DNA translocation, and in the case of short DNA strands, they cause deviations from a linear DNA conformation. When DNA is under tension, translocation is once again slower when aromatic residues are present in the constriction. However, the lack of flexibility in the DNA backbone provides a narrower window of opportunity for the DNA bases to be retained inside the pore via interaction with the aromatic residues, compared to more flexible strands. Consequently, there is more variability in translocation rates for strands under tension. DNA entry into the pores is correlated to pore width, but no such correlation between width and translocation rate is observed.

---

DNA sequencing using nanopores is now a well-established ‘next generation sequencing’ method^1^. Over the years there has been much attention devoted to optimizing the apertures within the membrane, with the aim of controlling conformation and translocation rate of DNA to improve the resolution of the devices^2–4^. The apertures are encased within nanoscale pores which are based on proteins or synthetic materials. Protein engineering of the *S. aureus* toxin α-hemolysin (a heptamer that forms a 14-stranded β-barrel in the membrane) followed by electrophysiology measurements revealed that basic residues introduced into the lumen of the pore of α-hemolysin slowed down the translocation of ssDNA^2^. Subsequent simulation studies of simplified models of the α-hemolysin pore region showed that the strong electrostatic interactions between the basic residues and the acidic phosphate groups of the DNA led to transient tethering of the DNA to the pore, which for short DNA strands resulted in undesirable DNA conformations that may impact on the accuracy of the read^5^. The approach of using model pores was then employed to study hydrophobic hour-glass-shaped pores. These pores were based on a 14-stranded β-barrel architecture and contained a central constriction formed by amino acid residues GAVLVAG^6^. It was shown that ssDNA was slow to enter these pores, however once inside it remained in a largely linear conformation. However, in the absence of the constriction region, the DNA was observed to coil up somewhat. Inspired by experimental work on engineered α-hemolysin which showed some advantages in having two constriction regions within a nanopore^4^, here we have designed a set of four nanopores, each with two constriction regions, to explore the impact of pore geometry and chemical nature of the constriction on DNA translocation rate and conformation. ssDNA translocation through these pores, under an applied electric field, is investigated for two scenarios; short (12 bases) DNA strands, and long DNA strands under tension to retain a linear conformation.

Our results show that tryptophan and phenylalanine residues inside the pore interact with the DNA to slow down DNA translocation under an applied electric field. This effect is observed both with short strands of DNA that coil/bend due to the interaction, and also longer DNA strands under tension that cannot alter their conformation to any significant extent.

## RESULTS AND DISCUSSION

### Model nanopores in an applied electric field

The 4 nanopores studied here mimic β-barrel proteins. Each barrel is composed of either 14 or 16 anti-parallel strands and contains 2 hydrophobic regions. The details of the pores are as follows: 14LLx2: 14 stranded barrel with 2 constrictions, each composed of 2 rings of LEU residues, 14Fx2: 14 stranded barrel with 2 constrictions, each composed of 1 ring of PHE residues, 16FFx2: 16 stranded barrel with 2 constrictions, each composed of 2 rings of PHE residues, and 16WWx2: 16 stranded barrel with 2 constrictions, each composed of 2 rings of TRP residues. For all pores the constriction nearest the entrance to the pore is termed constriction 1, while the constriction closest to the pore exit is termed constriction 2. Two lengths of DNA were used in this study; a 12-nucleotide strand of finite length and a 40-nucleotide strand bonded to itself through periodic boundaries, both have a poly A sequence. (Figure 1).

**Figure 1:**
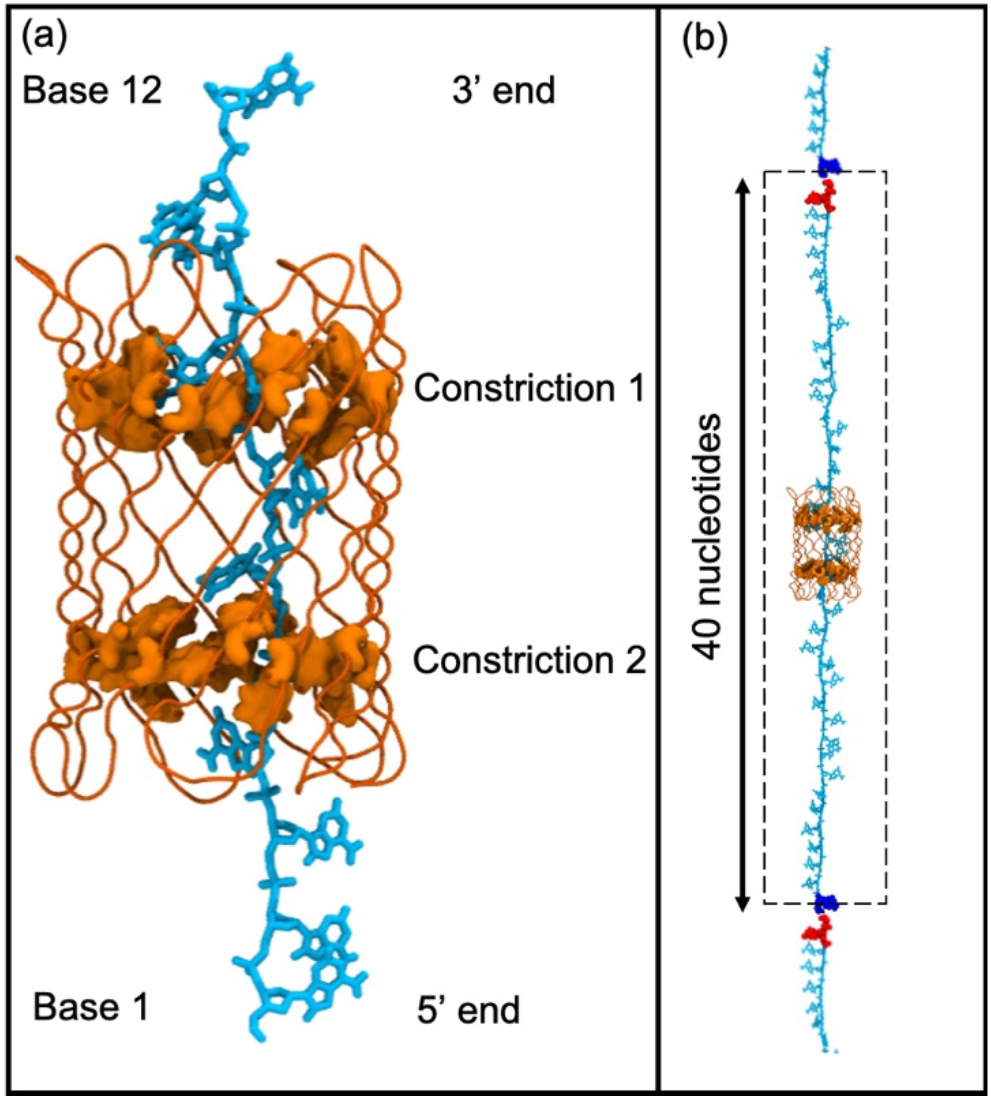
Panel (a) shows the naming conventions adopted throughout this study, and the initial position of the 12-nucleotide DNA (cyan) in the 14Fx2 pore (orange) for the translocation studies (the constriction residues are shown in surface representation). Panel (b) shows the continuous strand of DNA under tension. The terminal bases are colored blue and red to provide an illustration of the bonding across periodic boundaries (depicted by the dashed lines) to give a continuous strand of DNA.

We explored the behavior of the pores under an applied electric field. The dimensions of the pores fluctuate only by a small amount under an applied electric field of 0.15 V nm^−1^ under these conditions, due to side chain flexibility (Supporting Information). The radii of the constriction regions of the 14-stranded pores are ~0.47 nm and 0.58 nm in 14Fx2 and 14LLx2 respectively, with the 14Fx2 pore having a larger separation between the constrictions. The radii of the constriction regions in the 16-stranded pores are ~0.70 nm and 0.62 nm for 16FFx2 and 16WWx2 respectively.

### Entry of short DNA strands into hydrophobic pores under an applied electric field

The translocation of ssDNA through these pores was investigated. Two different initial distances from the pore were investigated; (1) the DNA terminal base (5’ end) is initially located at the entrance of the pore (2) the DNA is pre-threaded through the pore, such that bases 1-3 (where 1 is the terminal base at the 5’ end) are already outside the pore and only base 12 has not yet entered the pore. We have previously shown that the DNA will not enter 14-stranded hydrophobic pores (containing 1 constriction region), without an electric field, and rarely even with an applied electric field^6^. Here we observed that in all 8 simulations under an applied electric field of 0.15 V nm^−1^ in which the DNA is initially located at the mouth of the 14-stranded pores (4 × 14Fx2 and 4 × 14LLx2), DNA did not enter the pore at all. In contrast to this, in all 8 simulations of the 16-stranded pores (4 × 16FFx2 and 4 × 16WWx2), DNA only failed to translocate past constriction 1 once (movement of DNA as a function time for these simulations is provided in the Supporting Information). The simulations show that there is a clear barrier to DNA entry into the 14-stranded pores, but not the 16-stranded pore. This is likely to be related to the entropic penalty incurred upon entering a more confined geometry.

### Translocation of short DNA strands under an applied electric field

We next explored the simulations in which DNA had been pre-threaded through the pore. Six independent simulations were performed for each pore. The position of the center of mass of the 5’ terminal end base as a function of time was calculated to characterize the DNA translocation rate through the pores (Figure 2). In all 6 simulations of the 14LLx2 pore, we found the DNA to take between 4-17 ns to exit constriction 2, and in 2 of the simulations the DNA exited the pore completely (by 8 ns and 15 ns). Translocation was slower for the 14Fx2 pore, with the DNA remaining threaded through either both constriction regions (in 3 simulations) or constriction 2 only (in 3 simulations). For the 16-stranded pores, DNA did not exit constriction 2 in 5 and 4 of the 16WWx2 and 16FFx2 simulations respectively (Figure 2). For the former this was the case even when the simulations were extended to 40 ns.

**Figure 2:**
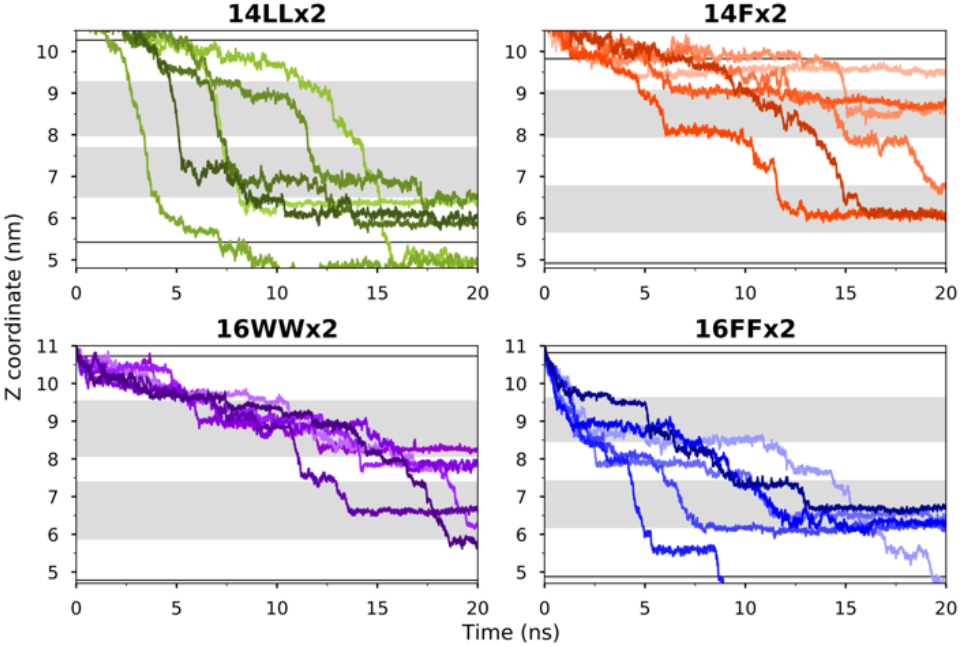
DNA translocation time in the short strand DNA simulations, measured as the Z coordinate of the center of mass of nucleotide 12 (3’-terminal end) as a function of time for all 6 simulations of each pore in which DNA is pre-threaded into the pore. The pore constriction regions are marked by dark grey bands, the solid lines represent the mouths of the pores.

We next sought to establish the origins of these differences in translocation rate by considering the DNA-pore interactions and the conformational behavior of the DNA strand. Cluster analysis was performed for a 6-nucleotide DNA segment by concatenating portions of the trajectories for each pore during which all bases of the segment are located inside the pore. The segments outside the pore are not included in the analysis as interactions with the mouth of the pore or the membrane may mask the effect of the interior of the pore; the latter is our focus here. The DNA end-to-end distances (phosphate to phosphate) for representative snapshots of each cluster are reported in Figure 3.

**Figure 3:**
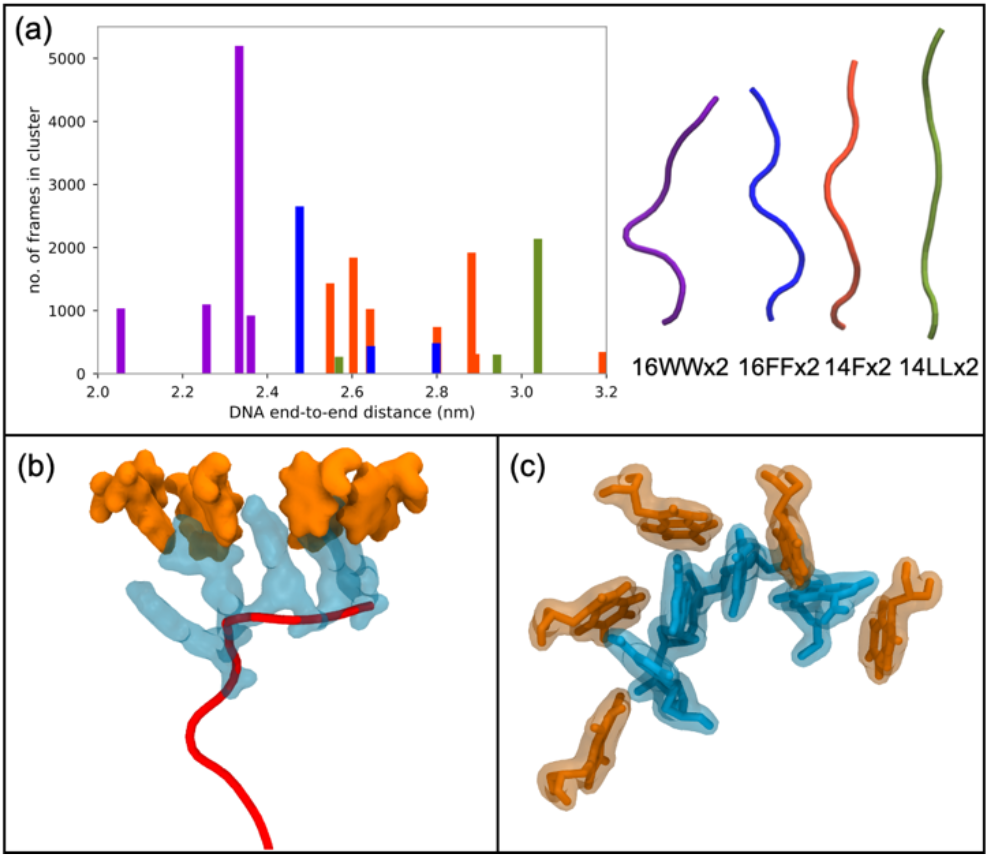
DNA conformation in the short strand DNA simulations. Panel (a) shows the distribution of end-to-end distances of DNA bases within each pore. The data is presented as the number of frames from the 6 trajectories for each pore that fall into each cluster. Representative backbone conformations are provided on the top right of the panel. Panel (b) shows 4 × DNA bases (cyan) interacting with TRP residues (orange) of the 16WWx2 pore. The concomitant distortion of the DNA backbone is evident (red). Panel (c) shows another view of these interactions in which the DNA bases are seen slotting into the gaps or pockets between the TRP residues.

We consider the 16-stranded barrels first. The end-to-end distance of DNA conformations inside these pores varies from 2.05-2.8 nm. The aromatic residues within the constrictions in the 16-stranded barrels show definite interactions with the DNA. These interactions have two consequences; they slow down the translocation of DNA and also cause it to deviate from a linear conformation. In general, these interactions arise as the aromatic sidechains form pockets within which the DNA bases can fit (Figure 3). In both pores, the interaction with the residues in the constriction regions causes the DNA to coil such that up to 4 bases are often interacting simultaneously with residues of a particular constriction. The end-to-end distance of major DNA conformations in the 16FFx2 pore varies from 2.5-2.8 nm, while this range is 2.05-2.35 nm for 16WWx2. In the 16WWx2 pore, in some cases when the bases at the 3’ terminus break free from constriction 1 they rapidly move towards constriction 2, as they are unhindered outside of the constrictions. However, due to the bases already within constriction 2 not being able to escape, there are occasionally up to 6 bases within this constriction leading to retention of the DNA inside the pore, and alterations in its conformation. Examples of the stepwise movement of the DNA between the two constriction regions is demonstrated by plotting the center of mass motion of each base as a function of time (Supporting Information). This effect is both less pronounced and less consistent for the 16FFx2 pore, leading to faster translocation times compared to 16WWx2 in general, although there is some variation. Within the 14LLx2 pore, the DNA is in a largely linear conformation throughout the time it spends inside the pore. Cluster analysis shows that the end-to-end distance of DNA conformations range from 2.55 to 3.05 nm (with ~ 80 % of DNA from all 6 simulations corresponding to the conformation with end-to-end distance of 3.05 nm). We note here that the DNA end-to-end distance is 3.4 nm at the beginning of the simulations. There is little interaction with the pore as the DNA moves through, despite the pore being narrow thus, there is no tethering effect. This results in relatively fast translocation rates while also maintaining the DNA in a largely linear conformation, such that there are only 1-2 bases inside the constriction region at any point. DNA within the 14Fx2 has clear interactions with the PHE residues that comprise the pore constrictions. The conformations of the DNA inside the 14Fx2 pore are generally more compact than those in the 14LLx2 pore, but comparable to those in 16FFx2. This is commensurate with the DNA translocation rate observed for this pore, which is slower than 14LLx2. The results here clearly indicate that the interaction with aromatic residues leads to less linear DNA structures and slower translocation rates.

### Translocation of long, tensioned DNA strands under an applied electric field

We next performed a set of simulations to explore if the pattern in DNA translocation rates are reproduced when the DNA conformation is forced to be more extended. A longer strand of polyA DNA composed of 40 nucleotides was bonded across periodic boundaries in the z dimension to give a continuous polymer threaded through the pores. The DNA was simulated in the NVT ensemble and, as the box size is kept constant, the DNA is essentially under tension and cannot coil/kink. The continuous nature of the DNA enables us to perform one long simulation (200 ns) for each pore and observe multiple DNA translocation events. Interestingly, once again we observe that the rate of translocation is the fastest through the 14LLx2 pore, and in general the DNA is retained for longer durations through the 3 pores with constriction sites formed by aromatic residues, when simulated with an applied field of 0.09 V nm^−1^. While DNA translocation is uniformly rapid through 14LLx2 (~40 ns for translocation of all 40 nucleotides) in these simulations, the rate is more varied through the pores with aromatic constrictions (Supporting Information). For example, the first full translocation of 40 nucleotides occurred within 40 ns through 14Fx2, but the second translocation event did not fully complete within the 40-200 ns timeframe of the remainder of the simulation. In particular, during the time interval ~ 42-145 ns, there is no appreciable translocation of ssDNA through this pore. Inspection of the trajectory revealed two neighboring PHE residues in constriction 1 that ‘trap’ one of the DNA bases through non-specific steric interactions, forming a sort of gate. When these residues rearranged such that the separation between their aromatic rings was > ~ 0.5 nm, the base was observed to move out of the constriction under the influence of the electric field. The correlation in the distance between the PHE residues forming the ‘gate’ and the DNA translocation rate is clear (Figure 4).

**Figure 4:**
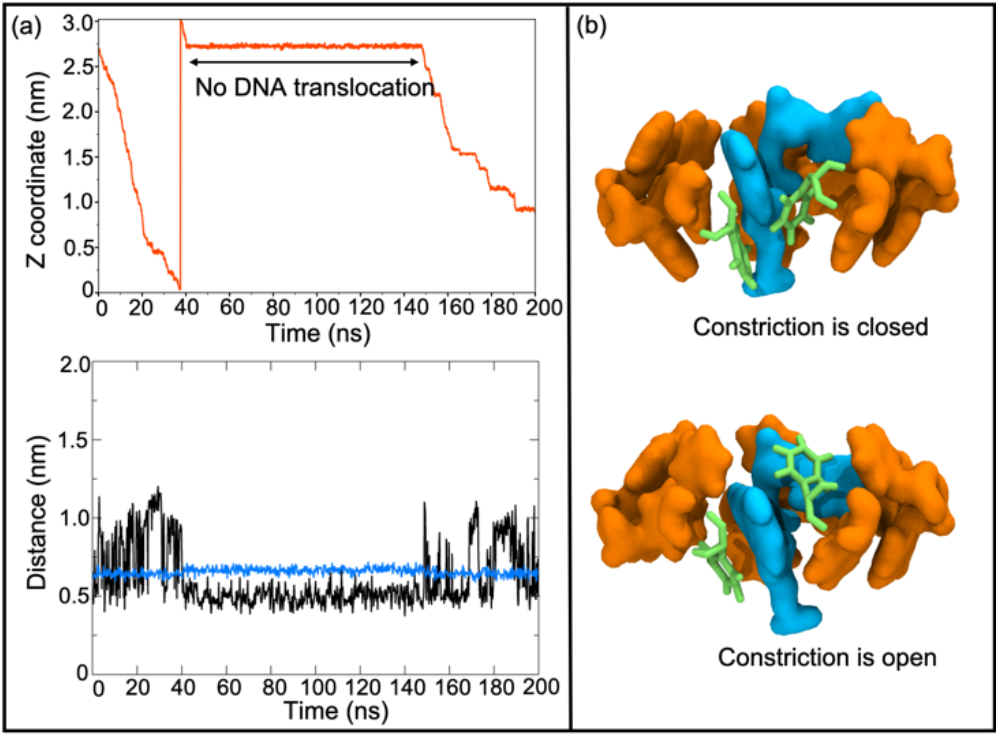
Panel (a) shows the center of mass movement of base 40 (3’ end) of the continuous DNA model as a function of time through the 14Fx2 pore (top). The bottom of panel (a) shows the distance between the aromatic rings of two neighboring PHE residues (black). The distance between the backbone Cα atoms of these residues (blue) is shown for comparison. Panel (b) provides a molecular view of the same PHE residues (green) in the context of other residues in constriction 1 (orange) and their interaction with two DNA nucleotides that occupy the constriction from time = 42-145 ns (cyan).

Similarly, for the 16-stranded pores, DNA retention times can be correlated with interactions with aromatic residues in the constriction. The aromatic residues are observed to form tight pockets around the DNA bases (Supporting Information), through non-specific interactions. Steps in the translocation correspond to bases moving out of a pocket, and others being retained either within the same constriction or the other one (Figure 5).

**Figure 5:**
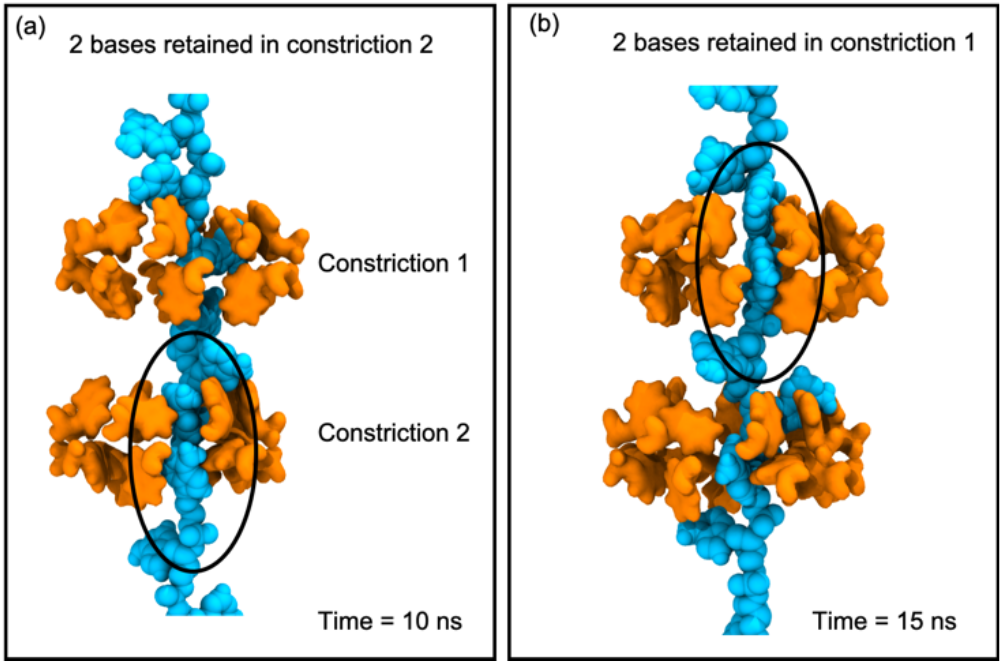
Panel (a) shows two bases caught in a pocket formed by TRP residues in constriction 2, and panel (b) shows the same system 5 ns later, now two bases are caught within a pocket in constriction 1. While there are none in the pockets of constriction 2 (1 base is present inside constriction 2, but this is a loose association, rather than a tight fit into a pocket). This leads to stepwise translocation of the ssDNA.

The faster translocation events observed for these pores (full translocation in ~ 40 ns), correlate with the timescales for full translocation through the 14LLx2 pore. Visual inspection of the trajectories reveals that in these cases of rapid translocation, the DNA is unable to move into a tight-fitting pocket. There are two factors that contribute to this; (a) the DNA is moving under the field and (b) it has very little conformational freedom in the backbone. Consequently, there is a chance that the DNA will not be retained within a pocket formed by neighboring aromatic residues due to the initial interaction not being optimal, and given it cannot alter its backbone conformation to optimize/improve the interaction, it is able to translocate rapidly past the constriction. Interestingly, the pores containing PHE residues in their constrictions retained tensioned ssDNA for longer periods than 16WWx2. We reason that this likely due to the greater flexibility of the PHE side chains compared to the bulkier TRP side chains (root mean square fluctuation data is provided in the Supporting Information), which enable the former to rapidly form a pocket around DNA bases before they can move through the pore.

## CONCLUSIONS

In conclusion, we find that DNA enters the 16-stranded pores readily, but not the 14-stranded pores under an applied electric field. Thus, DNA entry is strongly influenced by the pore width. Short strands of DNA move through 14LLx2 unhindered and in a linear/extended conformation. The presence of aromatic residues in the constriction slows down the DNA translocation rate and also leads to DNA deviating from a linear conformation. The slowest translocation rates are observed for 16WWx2. This pore is intermediate in width between the other two pores with aromatic constriction regions. Thus, the nature of the residues at the constriction region has a strong influence over DNA translocation rates for short strands, but there is no correlation between pore width and translocation rate for the pores studied here. For DNA strands under tension, once again the fastest translocation rates are for 14LLx2. Pores containing aromatic residues, in particular PHE, can retain the DNA via interactions with the bases for extended periods thereby slowing down translocation. However, if at least one base is not retained within a pocket formed of aromatic residues, the DNA translocation rate is the same as within 14LLx2. In terms of design principles for nanopores used within DNA sequencing devices, our results suggest pores engineered to include TRP for flexible DNA strands and PHE for DNA under tension can slow down translocation rates.

## MATERIALS AND METHODS

The hydrophobic pore models used in this work were those constructed using Modeller and PyMOL and were based on the pores first reported by Sansom and co-workers^7^. The ssDNA model used was a 12-nucleotide polyA strand, and was generated using the 3DNA package.^8^ The Cα atoms of the pores were restrained using a harmonic potential with force constant 1000 kJ mol^−1^ nm^2^.

Simulations were performed using GROMACS versions 5.1.4 and 2016, corresponding to updated package releases, and the GROMOS 53a6 forcefield with Berger lipid definitions.^9–11^ Pores were inserted into bilayers of 505 DPPC lipids, solvated using the SPC water model and ions added to a concentration of 1.0 M.^12^ Additional ions were added to neutralize systems prior to simulation. Positional restraints with force constant 1000 kJ mol^−1^ nm^2^ were applied to the phosphate moieties of DPPC lipids to prevent movement in the z dimension. Long-range electrostatics were treated using the Particle Mesh Ewald method with a short-range cutoff of 1.4 nm.^13^ The van der Waals interactions were truncated at 1.4 nm with long-range dispersion corrections applied to the energy and pressure. The temperature of the systems was maintained at 310 K using the v-rescale thermostat and a coupling constant of 0.1 ps for both short and continuous DNA systems. The continuous DNA was simulated in the NVT ensemble, whereas the short DNA was simulated in the NPT ensemble. Pressure was maintained at 1 bar semi-isotropically using the Parrinello-Rahman barostat and a time constant of 1 ps. ^14^ All bonds were constrained using the LINCS algorithm enabling a timestep of 2 fs.^15^ The electric field was imposed by a constant voltage drop across the simulation cell. Periodic boundary conditions were applied in three dimensions as in our own previous work on pores based on α-hemolysin and the work of other groups studying similar phenomena.^5, 16–18^ Analysis was conducted using GROMACS utilities and locally written code. Pore radius profiles for the pore models were obtained using HOLE.^19^ Molecular graphic images were produced using the Visual Molecular Dynamics, VMD package.^20^

## ASSOCIATED CONTENT

### Supporting Information

The Supporting Information is available free of charge on the ACS Publications website. PDF file containing a radius profile of the pore under an applied electric field, DNA end-to-end distances and plots of DNA movement through the pores.

## AUTHOR INFORMATION

### Funding Sources

No competing financial interests have been declared. PR and TH are funded by Oxford Nanopore Technologies. SK is funded by BBSRC, EPSRC, and The Leverhulme Trust.

## Acknowledgments

The authors thank Mark Sansom and Shanlin Rao for helpful discussions. The authors acknowledge the use of the IRIDIS High Performance Computing Facility, and associated support services at the University of Southampton and the use of the UK national supercomputer, ARCHER granted via the UK High-End Computing Consortium for Biomolecular Simulation, HECBioSim (http://hecbiosim.ac.uk), supported by EPSRC (grant no. EP/R029407/1), in the completion of this work.

## Table of Contents artwork

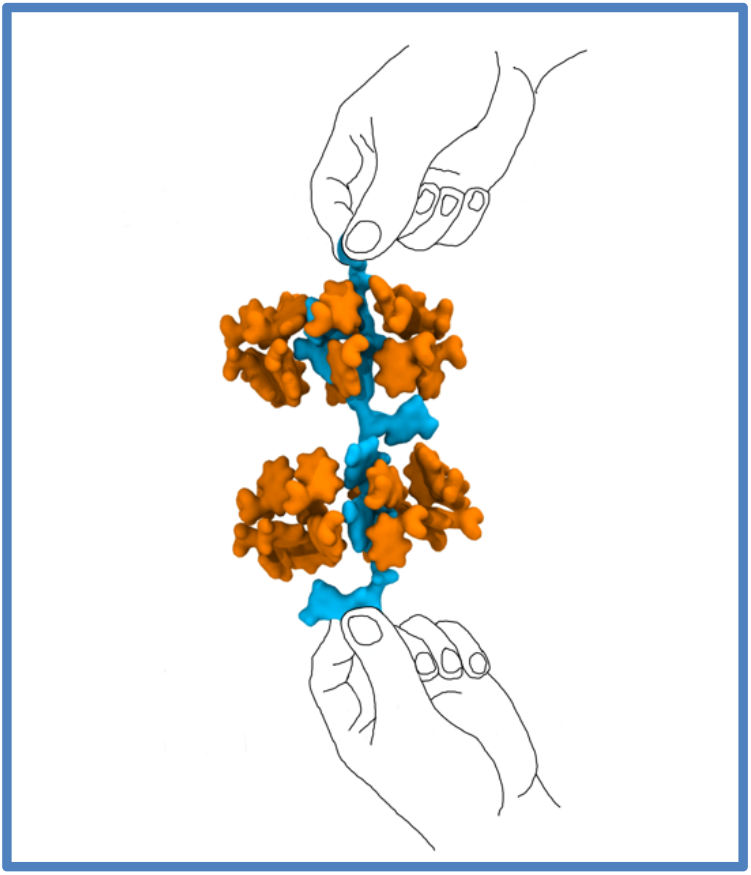

## Supplementary information

**Table 1.** A summary of all short strand simulations in which ssDNA was initially placed at the entrance to the pore

| Model pore | Final DNA location (number of simulations) |  |  |
| --- | --- | --- | --- |
|  | DNA above or in constriction 1 | DNA above or in constriction 2 | DNA at pore exit |
| 14LLx2 | 4 | 0 | 0 |
| 14Fx2 | 4 | 0 | 0 |
| 16FFx2 | 0 | 0 | 4 |
| 16WWx2 | 0 | 1 | 3 |

**Table 2.** A summary of all short strand simulations in which ssDNA was pre-threaded into the pore.

| Model pore | Final DNA location (number of simulations) |  |  |
| --- | --- | --- | --- |
|  | DNA above/ in constriction 1 | DNA above/ in constriction 2 | DNA in pore exit |
| 14LLx2 | 0 | 0 | 6 |
| 14Fx2 | 3 | 3 | 0 |
| 16FFx2 | 0 | 4 | 2 |
| 16WWx2 | 1 | 5 | 0 |

**SI Figure 1:**
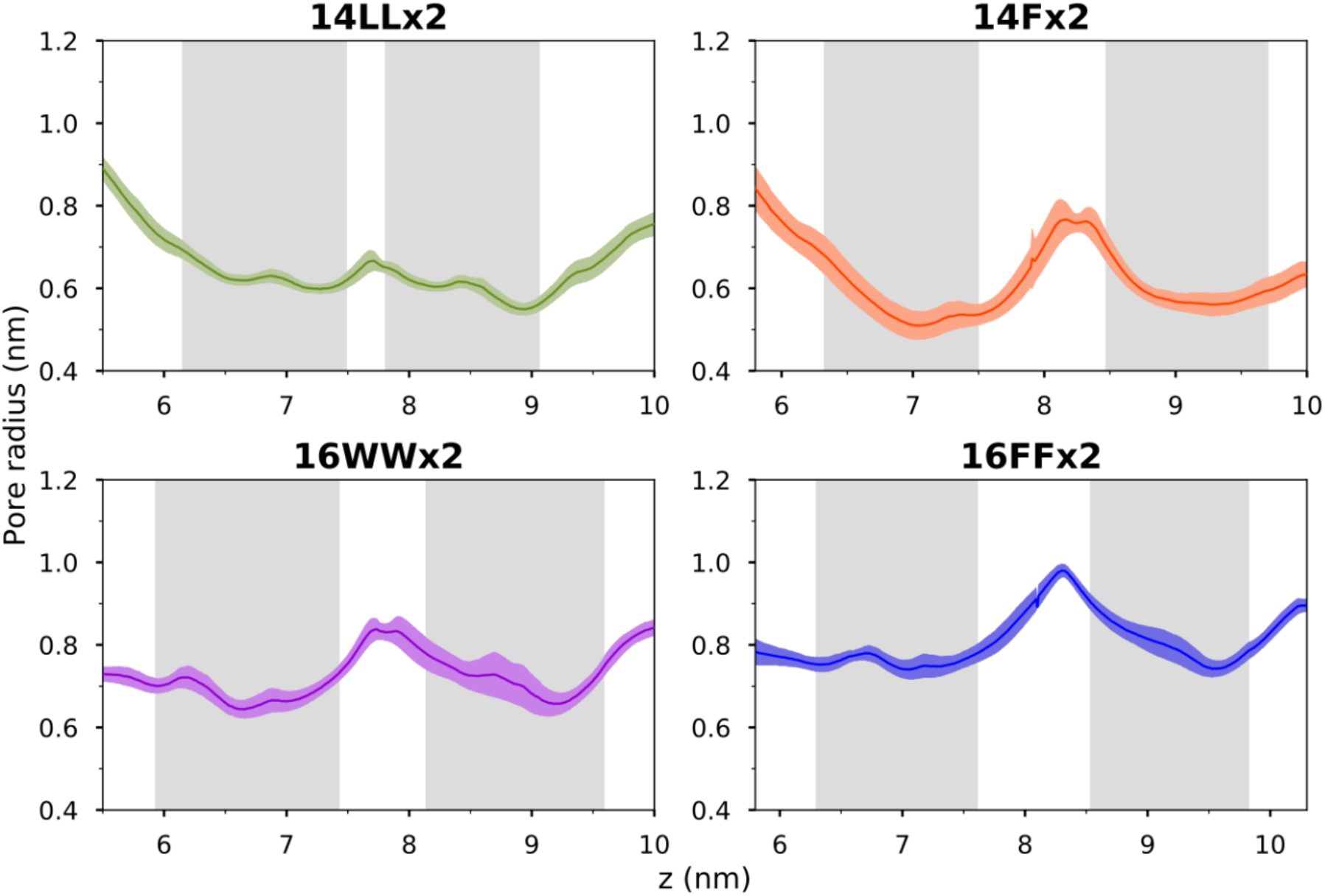
Average pore profiles for all four pores from simulations under an electric field of 0.15 Vnm^−1^, in the absence of DNA. Constriction regions are bounded by the two central vertical lines.

**SI Figure 2:**
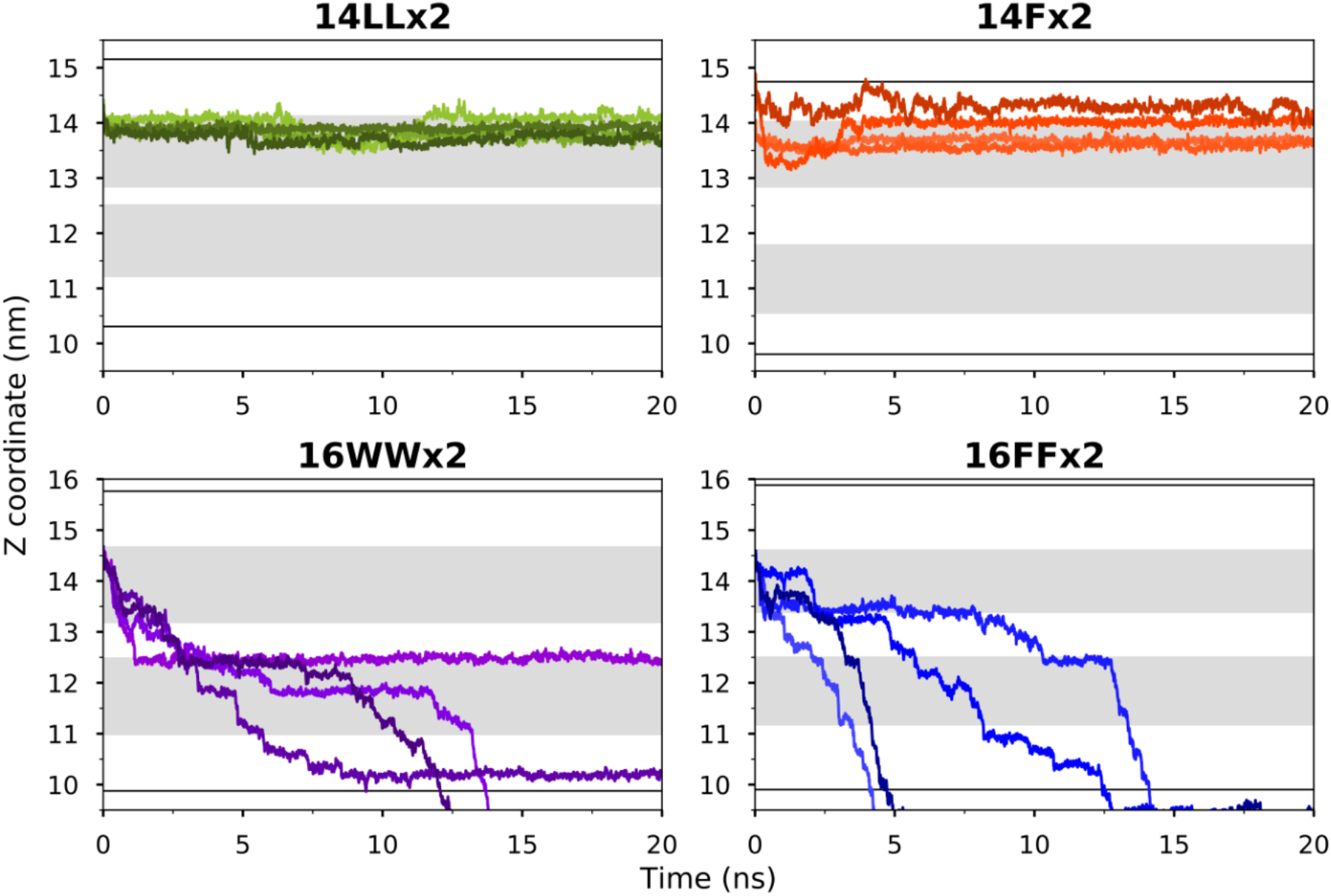
DNA translocation time in the short strand DNA simulations, measure as the Z coordinate of the center of mass of nucleotide 12 (3’-terminal end) as a function of time for all 6 simulations of each pore in which DNA is pre-threaded into the pore. The pore constriction regions are marked by dark grey bands, the solid lines represent the mouths of the pores.

**SI Figure 3:**
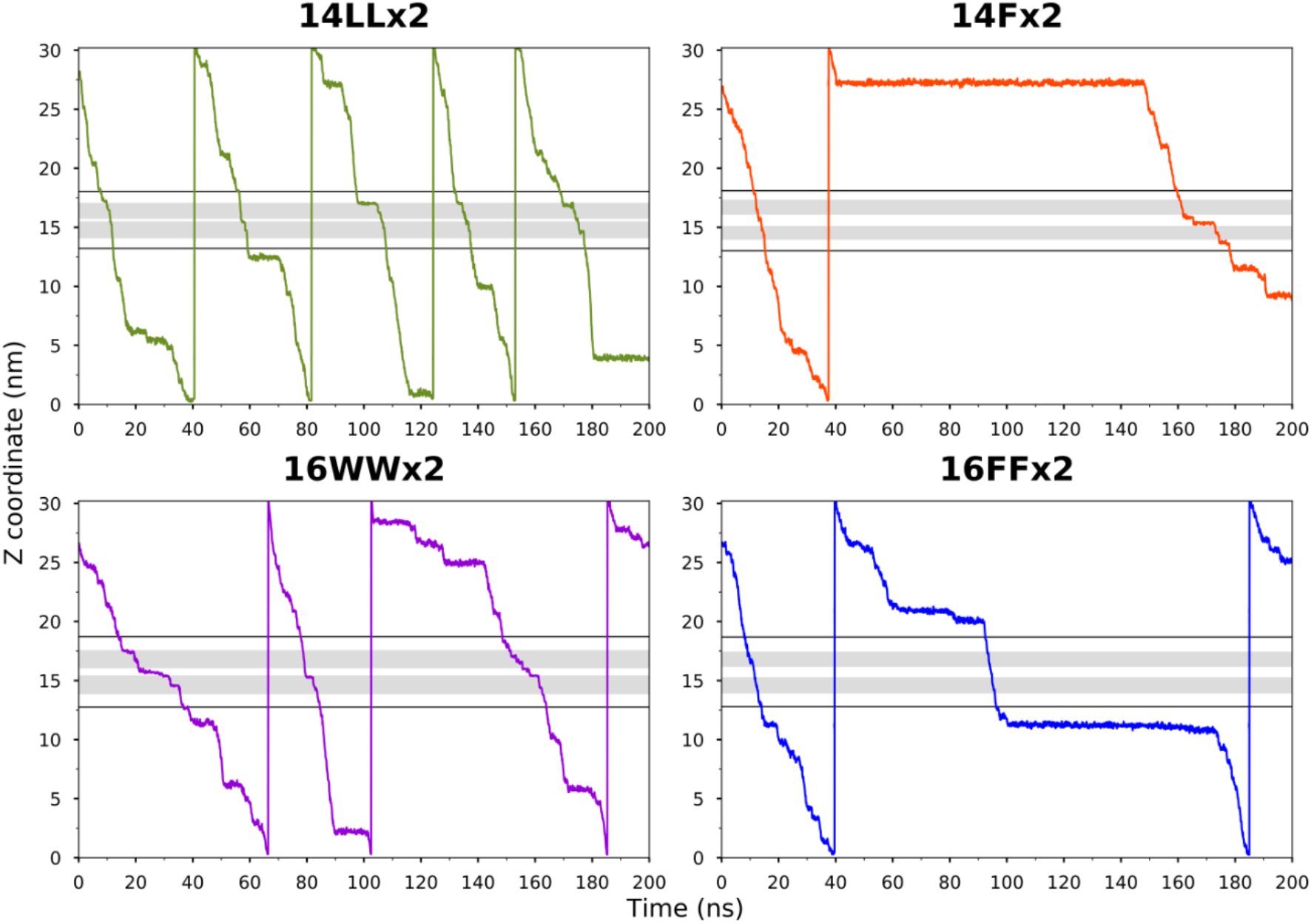
Translocation of continuous ssDNA under tension through all four pores. The center of mass of the base furthest from constriction 1 at the start of the simulations (3’) plotted against time. The shaded regions represent the pore constrictions and the solid black lines represent the mouths of the pores.

**SI Figure 4:**
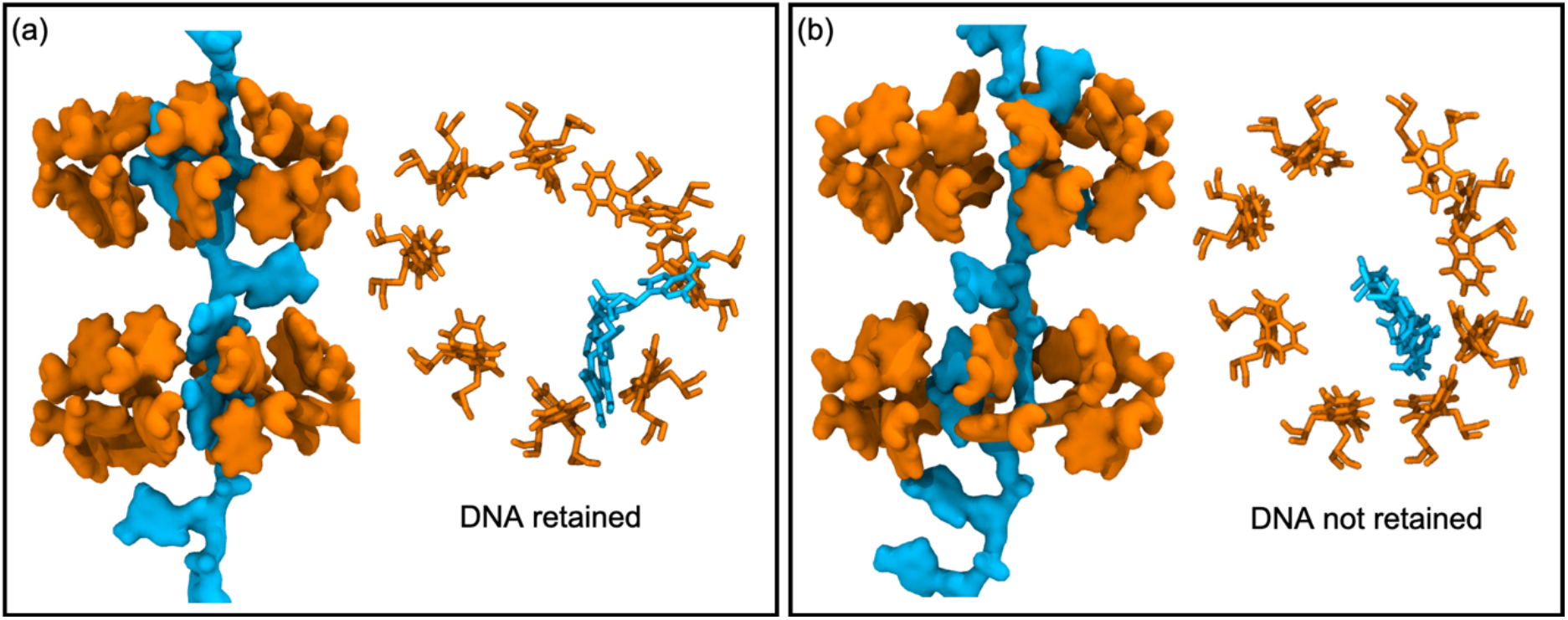
Two types of ssDNA translocation are observed through pores containing aromatic residues, rapid (~ 40 ns) and much longer. Panel (a) shows DNA bases (blue) retained within a pocket formed by TRP residues (orange) in constriction 2, top and side views this leads to tethering of the DNA to this region and thus slower translocation. Panel (b) shows a scenario in which the bases are unable to move into such a pocket leading to faster translocation.

**SI Figure 5:**
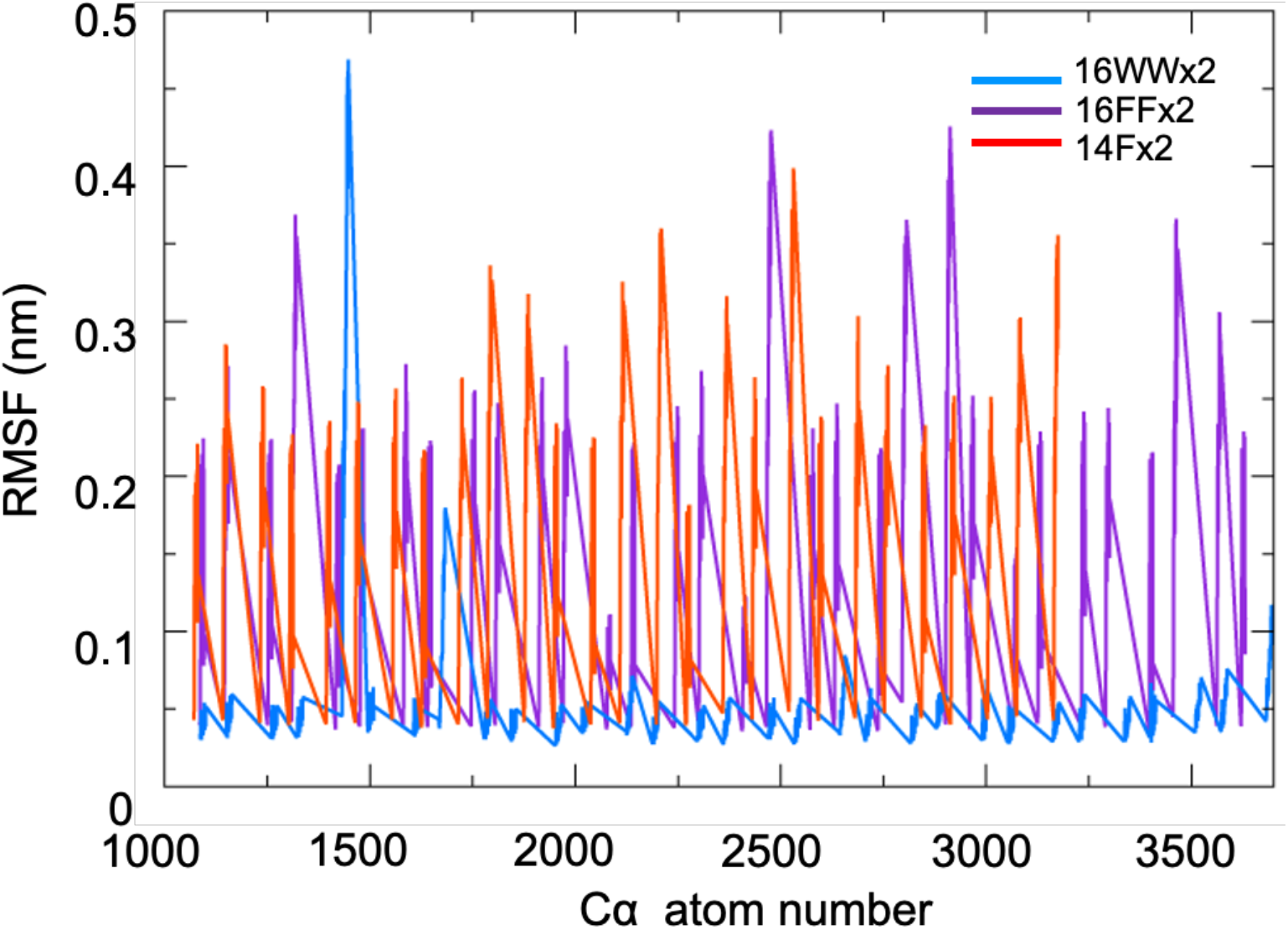
Root mean square fluctuations of the side chains of aromatic residues within the constriction regions of 16WWx2, 16FFx2 and 14Fx2. There is marked lower flexibility in the TRP side chains compared to the PHE sidechains.

## Notes

### Competing Interest Statement

The authors have declared no competing interest.

